# Cold exposure transiently increases resistance of *Arabidopsis thaliana* against the fungal pathogen *Botrytis cinerea*

**DOI:** 10.1101/2024.05.28.596154

**Authors:** Dominic Schütte, Abdulmalek Remmo, Margarete Baier, Thomas Griebel

## Abstract

A sudden cold exposure (4°C, 24 h) primes resistance of *Arabidopsis thaliana* against the virulent biotrophic pathogen *Pseudomonas syringae* pv. *tomato* DC3000 (*Pst*) for several days. This effect is mediated by chloroplast cold sensing and the activity of stromal and thylakoid-bound ascorbate peroxidases (sAPX/tAPX). In this study, we investigated the impact of such cold exposure on plant defence against the necrotrophic fungus *Botrytis cinerea*. Plant resistance was transiently enhanced if the *B. cinerea* infection occurred immediately after the cold exposure, but this cold-enhanced *B. cinerea* resistance was absent when the cold treatment and the infection were separated by 5 days at normal growth conditions. Plastid ascorbate peroxidases partially contributed to the transient cold-enhanced resistance against the necrotrophic fungus. In response to *B. cinerea*, the levels of reactive oxygen species (ROS) were significantly higher in cold-pretreated Arabidopsis leaves. Pathogen-triggered ROS levels varied in the absence of sAPX, highlighting the strong capacity for sAPX-dependent ROS regulation in the chloroplast stroma. The cold-enhanced resistance against *B. cinerea* was associated with cold-induced plant cell wall modifications, including sAPX-dependent callose formation and significant lignification in cold-treated Arabidopsis leaves.

**Funding:** This work was supported by the German Research Foundation (CRC973/C4) and the FU Berlin.

## INTRODUCTION

Abiotic factors, such as temperature, fluctuate strongly in most environments. In temperate regions, short temperature drops in spring are common and occur in irregular patterns. The frequency and risk of such late-spring frosts has increased in Europe and Asia within the last 60 years (Zohner et al., 2020). Sudden frost (< 0°C) often causes extracellular ice crystal formation, damages the plasma membrane, and reduces osmotically active water in the plant cells (Xin and Browse, 2000; Satyakam et al., 2022). Already chilling temperatures (0°C – 10°C) affect plant performance and growth. Such softer sudden cold exposures cause energy imbalances between the photosynthetic light reaction and the Calvin-Benson cycle and lead to the formation of reactive oxygen species (ROS) in chloroplasts (Huner et al., 1993; Ensminger et al., 2006), but also modify susceptibility of plants against pathogens. Several days after transition to 4°C, *Arabidopsis thaliana* (Col-0 accession) plants activate plant immune responses, such as salicylic acid production, and enhance immune gene expression even in the absence of pathogens (Kim et al., 2013; Kim et al., 2017). Daily repetitive 1.5 h cold treatments entrain and single 8-24 h lasting pre-exposures to 4°C prime plant resistance against subsequent infections with the bacterial pathogen *Pseudomonas syringae* pv. *tomato* DC3000 (*Pst*) (Singh et al., 2014; Wu et al., 2019; Griebel et al., 2022). In contrast to acclimation that describes plant adjustments to a sustained environmental change, entrainment results from repetitive and regular external cues, and priming is the outcome of an initial and single stress stimulus that influences plant response to a subsequent stress exposure (Hilker et al., 2016; Baier et al., 2019).

*Botrytis cinerea* is a necrotrophic pathogenic fungus and causes grey mould disease on plants (Williamson et al., 2007). It kills plant host cells as part of its infection strategy and can infect numerous plant species (Glazebrook, 2005; Williamson et al., 2007; Bi et al., 2023). Upon spore germination, *B. cinerea* forms specialized cells, so-called appressoria, to penetrate host epidermal tissues and cells (Bi et al., 2023). The fungal pathogen produces various cell wall-degrading enzymes, toxins, and oxalic acid to turn the host’s defence into susceptibility (Williamson et al., 2007; Nakajima and Akutsu, 2014). Plants employ an interconnected two-layered defence system against pathogen threats (Jones and Dangl, 2006; Jones et al., 2016; Ngou et al., 2021; Yuan et al., 2021): First, pattern recognition receptors (PRRs) anchored in plant membranes identify pathogen-associated molecular patterns (PAMPs) and initiate pattern-triggered immunity (PTI) (Zipfel, 2014; Albert et al., 2020). The PAMP chitin, for instance, is a major component of fungal cell walls and is detected by a receptor-like kinase known as Chitin Elicitor Receptor Kinase 1 (CERK1) (Miya et al., 2007). Second, intracellular receptors intercept pathogen virulence factors, so-called effectors, and activate effector-triggered immunity (ETI) (Jones et al., 2016; Lolle et al., 2020). This second layer, however, plays a negligible role against necrotrophic fungal pathogens and is rather important for defence against biotrophs (Liao et al., 2022).

Plant defence responses against *B. cinerea* include the generation of ROS, the biosynthesis of phytoalexins, such as camalexin, but also a fine-tuned activation of the plant hormones jasmonic acid (JA) and salicylic acid (SA), callose deposition, and cell wall modifications (Thomma et al., 1998; Ferrari et al., 2003; Veronese et al., 2006; Ferrari et al., 2007; Ramírez et al., 2011; Birkenbihl et al., 2012; Yang et al., 2018). Callose is a central component of papillae, which are locally and transiently formed plant cell wall modifications at the site of infection (Jacobs et al., 2003; Nishimura et al., 2003). The polymer lignin strengthens the plant cell walls and lignification is often enhanced in response to distinct biotic, but also abiotic stress treatments (Eynck et al., 2012; Cesarino, 2019; Lee et al., 2019; Nakamura et al., 2020; Ma, 2024).

ROS are important sub- and intra-cellular signalling molecules but also mediate cell-to-cell signalling (Miller et al., 2009; Ugalde et al., 2021; Peláez-Vico et al., 2024). In photosynthesis, ROS generation by the light-driven electron transport is unavoidable (Smirnoff and Arnaud, 2019; Foyer and Hanke, 2022). Plants neutralize these ROS by a highly efficient chloroplast antioxidant system. Tightly functionally interacting with superoxide dismutases, chloroplast ascorbate peroxidases (APX) utilize ascorbate as an electron donor to detoxify H_2_O_2_ (Groden and Beck, 1978). Thylakoid-bound APX (tAPX) is part of a first layer of protection and scavenges photosynthesis-related H_2_O_2_ directly at the thylakoid membrane (Asada, 1999; Jardim-Messeder et al., 2022). Stromal APX (sAPX) provides downstream antioxidant protection in the plastid stroma (Asada, 1999; Jardim-Messeder et al., 2022). Increasing evidence suggests that both plastid APX have additional functions in chloroplast energy metabolism and signalling (Kangasjarvi et al., 2008; Maruta et al., 2010; van Buer et al., 2016; Maruta et al., 2016; van Buer et al., 2019; Seiml-Buchinger et al., 2022). tAPX specifically regulates cold priming-mediated repression of core stress-responsive genes during a subsequent cold exposure (van Buer et al., 2019). In response to the initial cold, tAPX promotes the suppression of the chloroplast NADPH dehydrogenase subunits and alters chloroplast-to-nucleus stress signalling (Seiml-Buchinger et al., 2022). While tAPX transcripts decrease in the cold and slowly rise again during the post-cold phase, sAPX transcripts are upregulated in the cold and are quickly reset at normal growth temperatures (van Buer et al., 2016; Griebel et al., 2022). Previous work showed that both plastid APX contribute to cold priming-enhanced resistance against the bacterial pathogen *Pst* (Griebel et al., 2022). Even in the immune-compromised null mutant *enhanced disease susceptibility 1-2* (*eds1-2*), which is competent to establish cold priming-mediated *Pst* resistance, lack of sAPX abolishes cold priming-enhanced resistance against *Pst* (Griebel et al., 2022; Schütte et al., 2024).

Here, we investigated whether a single cold exposure (4°C, 24 h) impacts not only resistance of Arabidopsis against the virulent bacterial pathogen *Pst* but also against the necrotroph fungus *B. cinerea*. In addition, we analysed the role of plastid APX for cold exposure-enhanced fungal resistance and clarified their contribution for (post-)cold and *B. cinerea*-triggered ROS and callose deposition.

## RESULTS

### Cold pretreatment of Arabidopsis transiently enhances resistance against *Botrytis cinerea*

A 24 h cold exposure (4°C) of Arabidopsis Col-0 plants results in enhanced resistance against the bacterial hemibiotrophic pathogen *Pseudomonas syringae* pv. *tomato* (*Pst*) if the inoculation occurs immediately after the cold treatment, but also 5 days later (Griebel et al., 2022). Here, we studied the impact of such a prior cold exposure (4°C, 24 h) on plant resistance against the necrotrophic fungal pathogen *Botrytis cinerea*. Two different experimental setups were compared (Fig. 1A): (i) The first one consisted of the cold pretreatment (4°C, 24 h) and an immediately (2 h) subsequent infection (CT), (ii) the second included a stress-free period of 5 days at regular growth conditions (day/night: 20°C/18°C) between the initial cold exposure and the inoculation, and was therefore designated as cold priming (CP) setup. While the infection in the CT setup might be affected by the post-cold deacclimation, the infection 5 days after the cold exposure requires a priming memory, because cold-responsive genes and metabolites, such as soluble sugars and proline, are already reset after three days of deacclimation (Byun et al., 2014; Zuther et al., 2015; Griebel et al., 2022).

**Figure 1.**
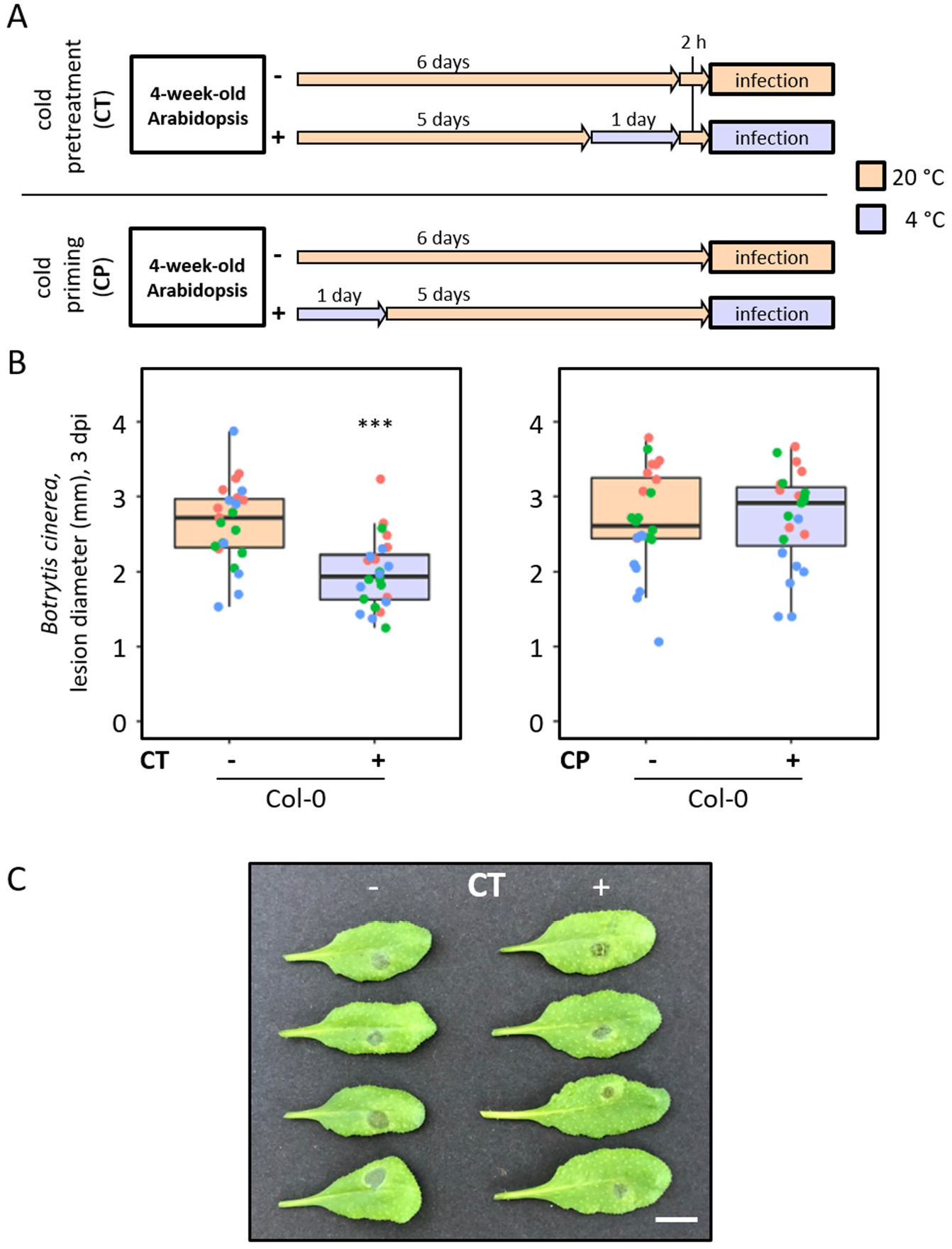
Impact of a prior cold exposure on resistance against *Botrytis cinerea* in *Arabidopsis thaliana*. **(A)** 5-week-old *Arabidopsis thaliana* (accession Col-0) plants were cold-pretreated (CT; 4°C, 24 h) and drop-inoculated (5 x 10^4^ spores ml^-1^) with *Botrytis cinerea* (*B*.*c*.) after 2 h at 20°C. For the cold priming setup (CP), 4-week-old Col-0 plants were cold-exposed (4°C, 24 h). Plants were drop-inoculated (5 x 10^4^ *B*.*c*. spores ml^-1^) after a memory phase of 5 days at normal growth conditions. **(B)** Lesion diameters of *B. c*.*-*infected plants were measured 3 days post inoculation (dpi) after preceding CT-treatment (left panel) or PT-treatment (right panel). Data points are shown from three independent experiments indicated by different colors (n= 23-24). Box plots show the median (central line) and asterisks denote statistically significant differences (*P* ≤ 0.05 (*), 0.01 (**), 0.001 (***), two-tailed *t-test*). **(C)** Representative picture of cold-pretreated (CT, +) and control (CT, -) Col-0 leaves with *B. c*. lesions at 3 dpi. Scale bar indicates 1 cm.

To study the impact of such prior cold treatments on resistance against *B. cinerea*, we analysed plant lesion sizes 3 days after drop inoculations with *B. cinerea* spores immediately and 5 days after the 24 h lasting cold exposure (Fig. 1B,C). Arabidopsis Col-0 exhibited significant smaller lesion diameters compared to naïve control plants, if the cold exposure immediately preceded the inoculation (Fig. 1B,C; CT). By contrast, lesions diameters in the CP group, with a 5-day memory phase after the cold, were similar to those in the control group (Fig. 1C). This indicates that a directly preceding cold exposure enhances the resistance of Col-0 against *B. cinerea*, but the effect is not sufficiently long-lasting to restrict fungal growth several days later.

### Plastid ascorbate peroxidases contribute to cold-enhanced resistance against *Botrytis cinerea*

Recently, we showed that cold-enhanced resistance against the bacterial pathogen *Pst* requires the plastid ascorbate peroxidases sAPX and tAPX (Griebel et al., 2022; Schütte et al., 2024). We repeated the experiment with *B. cinerea* including previously used *sapx* and *tapx* knockout lines (Kangasjarvi et al., 2008). In addition, we reduced the experimental set-up to the immediate cold pretreatment as this was effective in enhancing resistance against *B. cinerea* (Fig. 1). Again, cold-pretreated Col-0 plants showed significantly smaller *B. cinerea* lesions compared to the control group at 3 days post inoculation (dpi) (Fig. 2). In contrast, the difference in lesion sizes between cold-pretreated and control *sapx* and *tapx* plants was less pronounced and remained at a non-significant level (Fig. 2). This demonstrates that the availability of functional plastid peroxidases partially contributes to cold-enhanced resistance against the necrotrophic pathogen *B. cinerea*.

**Figure 2.**
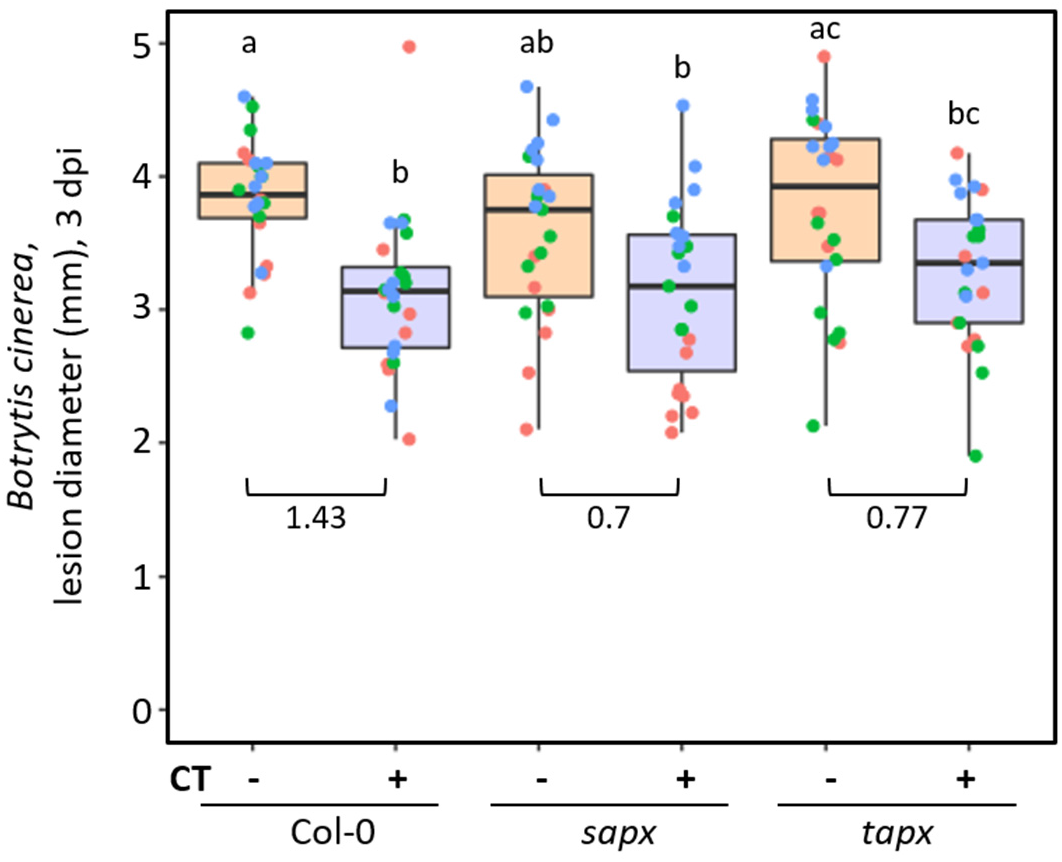
Impact of a cold pretreatment (CT; 24 h at 4°C) on resistance against *Botrytis cinerea* (*B*.*c*.) in Col-0 and in knockout lines of *stromal ascorbate peroxidase* (*sapx*) and *thylakoid ascorbate peroxidase* (*tapx*). Leaves of cold-pretreated (CT, +) and control (CT, -) plants were drop-inoculated (5 x 10^4^ *B*.*c*. spores ml^-1^) and the lesion diameter was measured 3 days post inoculation (dpi).). Data points are shown from three independent experiments indicated by different colours (n = 23-24). Box plots show the median (central line) and letters denote statistically significant differences (Tukey-HSD; *P* ≤ 0.05). Numbers between two boxes show the effect size between two means according to Cohen’s d.

Next, we tested transcript levels of selected *B. cinerea*-triggered defence genes for cold- or APX-related differential activation patterns. *Pathogenesis-related 1* (*PR1*) is responsive to SA, while *Plant Defensin 1*.*2* (*PDF1*.*2*) and *Pathogenesis-related 4* (*PR4*) belong to the group of pathogen-triggered genes that depend on JA signalling (Thomma et al., 1998). *Phytoalexin-deficient 3* (*PAD3*) encodes the cytochrome P450 enzyme CYP71B15 which catalysis the final biosynthetic step of the phytoalexin camalexin (Zhou et al., 1999; Schuhegger et al., 2006). A prior cold treatment did not result in altered *PR1, PDF1*.*2, PR4* and *PAD3* transcript levels 2 hours after the cold exposure (0 dpi) in the plant lines Col-0, *sapx*, and *tapx* (Fig. 3). Consequently, the cold pre-treatment had no impact on the level of the analysed transcripts prior to *B. cinerea* infection. (Fig. 3). At 1 dpi and 2 dpi after inoculation with *B. cinerea*, relative expression of *PR1, PDF1*.*2a, PR4*, and *PAD3* was increased. However, the prior cold treatment did not significantly impact *B. cinerea*-triggered transcript levels in Col-0, *sapx*, and *tapx* (Fig. 3). Since these genes are strongly connected with SA, JA, or camalexin pathways, it indicates that cold-enhanced resistance against *B. cinerea* does not correlate with altered SA and JA signalling or increased biosynthesis of camalexin.

**Figure 3.**
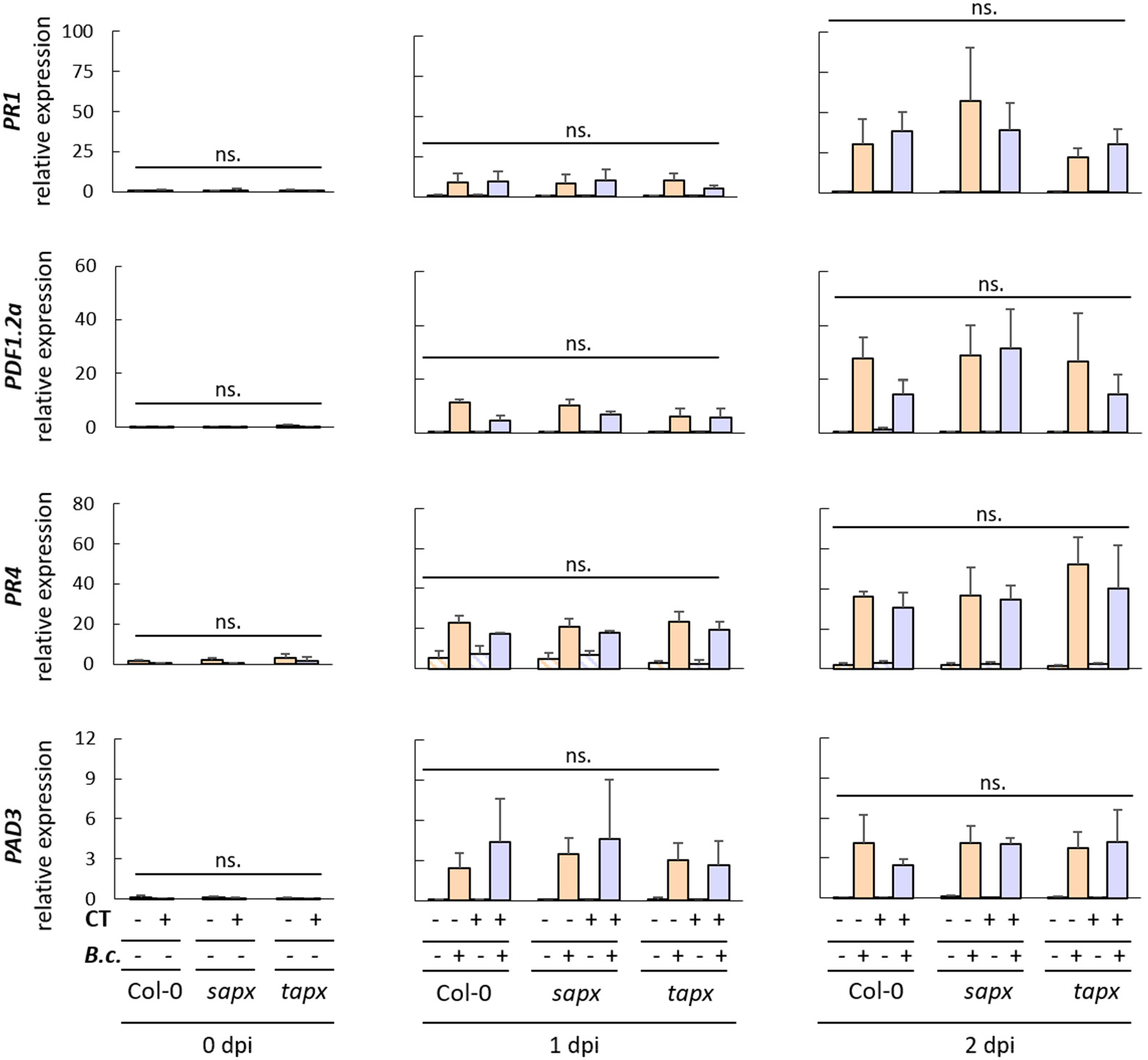
*Botrytis cinerea* (*B*.*c*.)-triggered transcripts of *PR1, PDF1*.*2a, PR4*, and *PAD3* in Col-0, *stromal ascorbate peroxidase* knockout (*sapx*) and *thylakoid ascorbate peroxidase* knockout (*tapx*) lines. Transcript levels in leaves were analysed after cold pretreatment (CT +; 4°C, 24 h) and subsequent *B*.*c*. spray infection (+; 2 x 10^5^ *B*.*c*. spores ml^-1^) at 0 days post inoculation (dpi), 1 dpi, and 2 dpi. Mock spraying (-; Vogel buffer) was used as infection treatment control. Transcript levels were determined by quantitative real-time PCR and were calculated as relative expression to the geometric mean of the reference genes *YLS8* and *RHIP1*. Bars show the mean of 3 independent experiments (n = 3) and the standard deviation. For mock treatments, only 2 independent experiments were performed and analysed (n = 2). No significant differences were observed for the comparison between the *B*.*c*.-treated samples (not significant, ns.; Fisher`s LSD test, *P* ≤ 0.05).

### *B. cinerea-*triggered ROS generation is enhanced in cold-pretreated Col-0 and in the absence of sAPX

sAPX and tAPX have central functions in scavenging photosynthesis-related H_2_O_2_ in the plastids (Asada, 1999). At the end of a 10 or 24 h cold exposure to 4 °C, H_2_O_2_ is significantly enhanced in Arabidopsis Col-0 leaves (van Buer et al., 2016; Wu et al., 2019). We analysed H_2_O_2_ levels by staining leaves with 3, 3′-diaminobenzidine (DAB) (Bittner et al., 2020). At the time point of inoculation (2 hours after the cold; 0 dpi), we could not detect increased H_2_O_2_ in cold-pretreated or naïve Col-0, *sapx*, and *tapx* (Fig. 4A). This is consistent with a rapid downregulation of cold-triggered ROS in Arabidopsis under optimal growth conditions given that enhanced ROS levels are detectable directly after the cold pretreatment (van Buer et al., 2016). 1 dpi after *B. cinerea* treatment, H_2_O_2_ remained at a low, not detectable level in Col-0, but it was significantly increased, if the plants where cold-treated before (Fig. 4A,B). Similarly, the *tapx* line showed enhanced H_2_O_2_ accumulation after *B. cinerea* infection (1 dpi) in prior cold-exposed leaves (Fig. 4A,B). In contrast to Col-0 and *tapx*, we detected strong *B. cinerea*-induced H_2_O_2_ accumulation in non-cold-treated *sapx* at 1 dpi, and the prior cold exposure even resulted in a weaker pathogen-triggered ROS accumulation in *sapx* (Fig. 4A,B).

**Figure 4.**
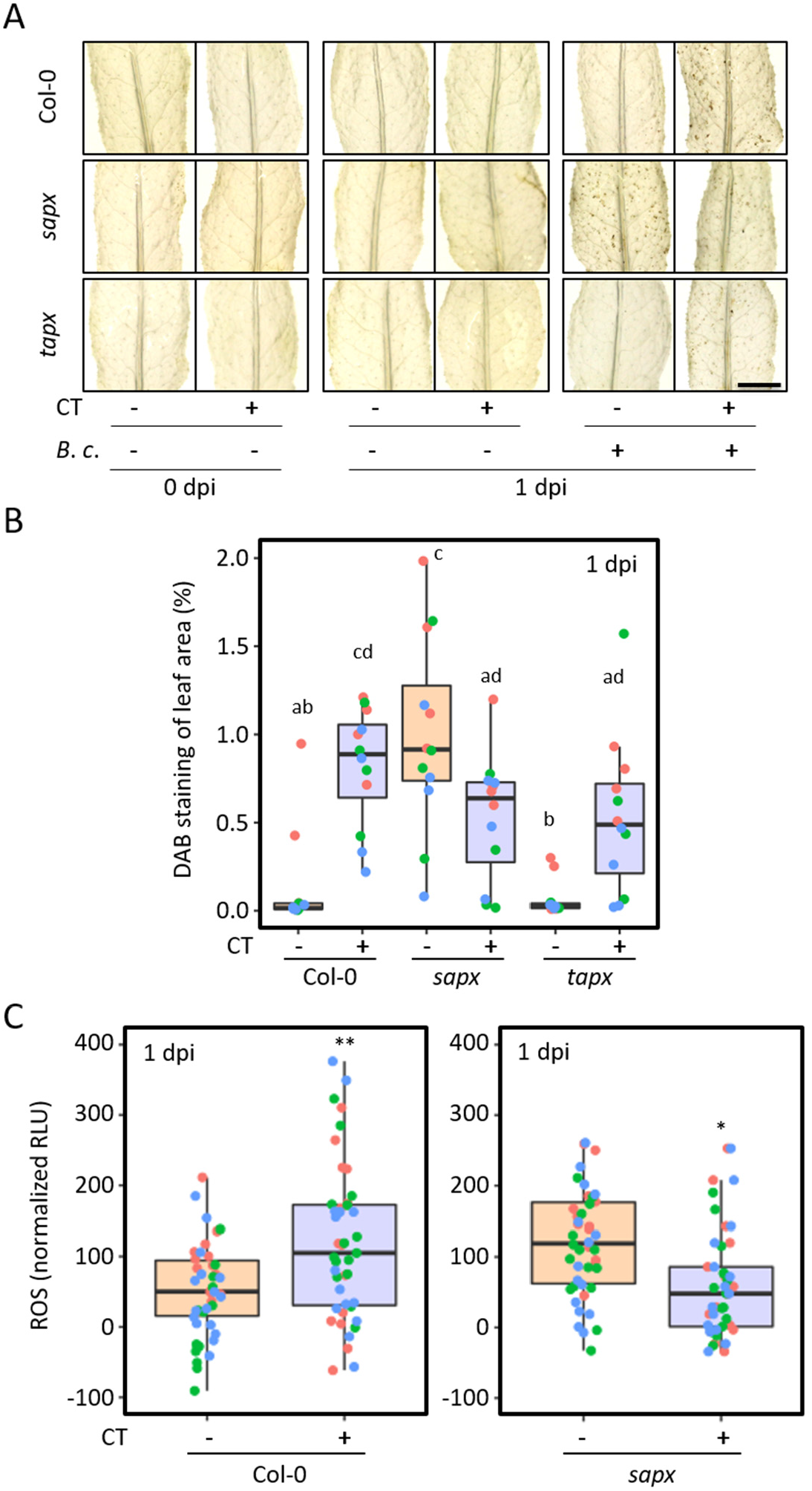
Reactive oxygen species (ROS) after cold pretreatment (CT +; 24 h at 4°C) and *Botrytis cinerea* spray infection (2 x 10^5^ spores ml^-1^ (+), Vogel buffer (-)) in leaves of Col-0, *stromal ascorbate peroxidase* knockout (*sapx*) and *thylakoid ascorbate peroxidase* knockout (*tapx*) lines. **(A)** Hydrogen peroxide accumulation at 0 days post infection (dpi) and 1 dpi visualized with DAB staining. Representative pictures are shown. Scale bar indicates 0.5 cm. **(B)** Quantification of DAB-stained H_2_O_2_. Pictures from (A) were analysed for the percentage of DAB-stained area of the leaf area. For each treatment and genotype 12 leaves from 3 independent experiments were analysed. Data points from independent experiments are shown in different colours. Box plots show the median (central line) and different letters denote statistically significant differences (Tukey-HSD: *P* ≤ 0.05). **(C)** Quantification of *B*.*c*.-triggered H_2_O_2_ with the luminol assay 1 dpi in Col-0 and *sapx*. Data points show the total amount of measured relative light units (RLU) counted within 1 hour relative to the non-infected control samples Data points from 3 independent experiments are shown in different colours (n = 46-48). Box plots show the median (central line) and asterisks denote statistically significant differences (two-tailed *t-test, P* ≤ 0.05 (*), 0.01 (**), 0.001 (***)).

To prove our results for Col-0 and *sapx* with an independent method, we used the luminol-based leaf disc assay for H_2_O_2_/ROS quantification (Bisceglia et al., 2015) and measured luminescence within a time frame of 1 h at 1 dpi. Although the differences were overall less pronounced, we observed the same trend as from the DAB staining (Fig. 4C): (i) *B. cinerea*-triggered ROS production was stronger in cold-pretreated Col-0 than in Col-0 that were constantly grown at normal growth conditions. (ii) Without a prior cold, *B. cinerea-*triggered ROS signals were significantly higher in *sapx* compared to Col-0 (p ≤ 0.001, t-test). (iii) Cold-pretreated *sapx* showed lower *B. cinerea*-triggered ROS levels than not cold-treated *sapx* plants suggesting that the antioxidant system overcompensates for the lack of sAPX.

### Cold-induced callose deposition requires sAPX

Callose, a β-1,3-glycan polymer, is transiently synthesized between the plasma membrane and the cell wall and contributes to the physical barrier (papillae) against pathogen attacks (Nishimura et al., 2003; Zavaliev et al., 2011; Ellinger and Voigt, 2014; Schneider et al., 2016). Callose biosynthesis and degradation in the neck region of plasmodesmata restricts their permeability and the transport of signalling compounds for intercellular communication (Zavaliev et al., 2011; German et al., 2023). In response to pathogens, callose deposition is initiated following detection of conserved pathogen-associated molecular patterns (Gómez-Gómez et al., 1999; Iriti and Faoro, 2009; Luna et al., 2011). However, callose can also be triggered by abiotic changes, such as cold (Wu et al., 2019). In some cases, increased callose accumulation correlates with enhanced resistance against *B. cinerea* in *Arabidopsis* (Nie et al., 2017; Sanmartín et al., 2020). We analysed callose deposition 2 h after the 24 h cold exposure (0 dpi) and in response to *B. cinerea* infection at 1 dpi. At 0 dpi, callose deposition in Col-0, but not in *sapx*, was significantly enhanced in cold-pretreated plants (Fig. 5 A,B). We could not anymore detect the cold-responsive callose in the non-infected Col-0 at 1 dpi. After infection with *B. cinerea* (Fig. 5 A,B), an increased callose deposition was observed in Col-0 and *sapx*. However, the number of *B. cinerea*-triggered callose spots was not affected by the cold treatment. We concluded that sAPX-promoted cold-triggered callose deposition correlates with enhanced resistance against *B. cinerea*.

**Figure 5.**
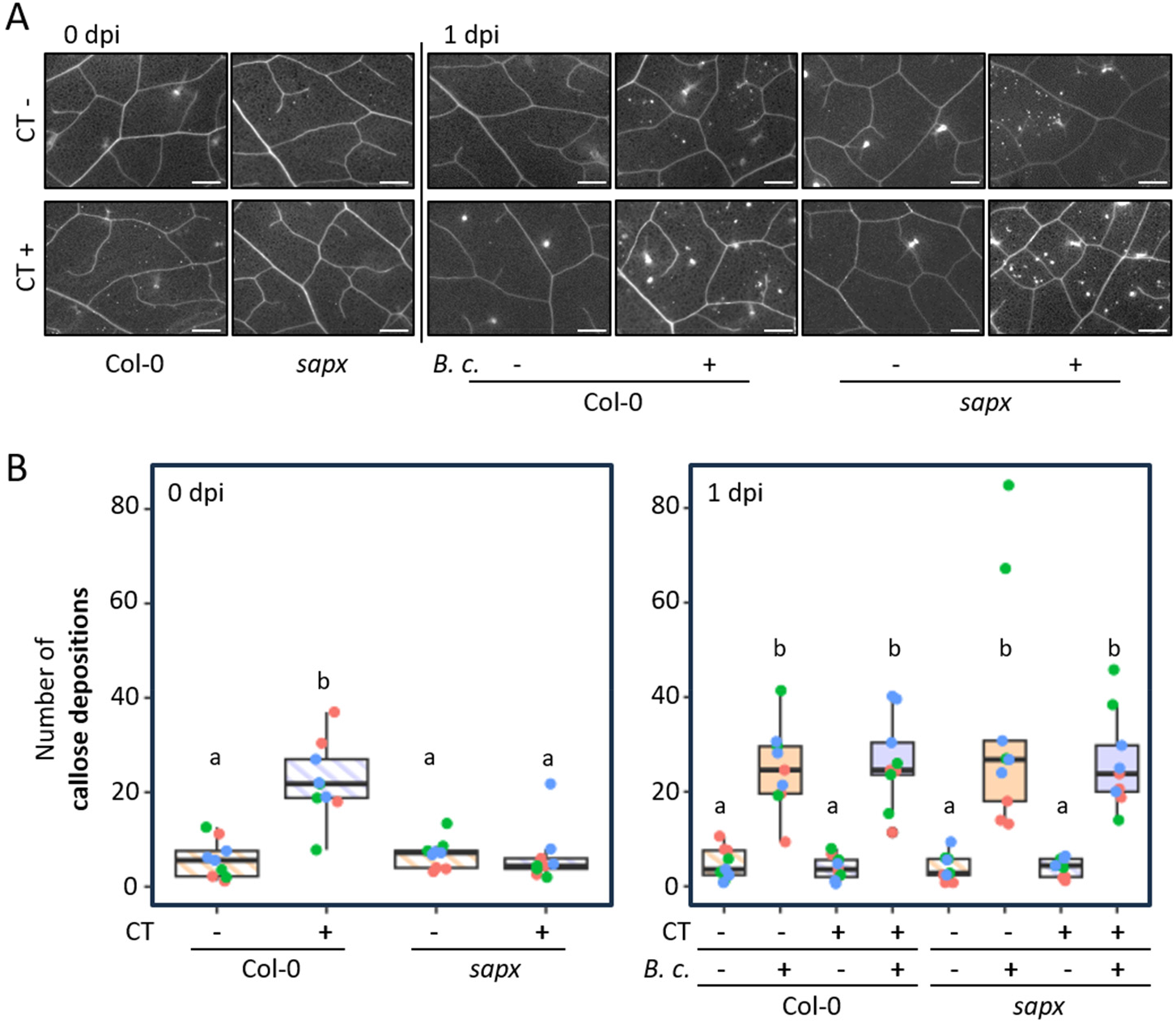
Callose formation after cold treatment (CT +; 4°C, 24 h) and subsequent *Botrytis cinerea* spray inoculation in Col-0 and *stromal ascorbate peroxidase* knockout (*sapx*). **(A)** Pictures of aniline blue-stained callose in Arabidopsis leaves immediately after CT (+, 0 days post inoculation (dpi)) and 1 dpi after subsequent *B. cinerea* infection (+) or mock infection (-) at 1 dpi. Scale bar = 500 μm. **(B)** Quantification of the callose depositions from (A). Each data point (biological replicate) represents the mean of 5 randomly analysed leaf areas (7 mm^2^). Counts from three independent experiments (n = 9) are shown and indicated by different colors. Box plots show the median (central line) and different letters denote statistically significant differences and different letters denote statistically significant differences (Tukey-HSD, *P* ≤ 0.05).

### Cold exposure enhances lignin content in cell walls

To analyse lignification of the cell wall in response to cold and *Botrytis cinerea*, we measured the lignin contents of extracted cell wall residues (CWR) using the acetyl bromide (AcBr) method (Moreira-Vilar et al., 2014; Chezem et al., 2017) in Col-0 and *sapx* leaves after prior cold exposure and 2 days after additional *B. cinerea* infection. The lignin content in the leaves of Col-0 significantly increased in response to cold (Fig. 6). Even at 2 dpi, the lignin ratio was still enhanced in cold-pretreated Col-0 samples, but not further altered in the *B. cinerea*-inoculated samples. This demonstrated that solely cold exposure but not fungal invasion was responsible for the significantly higher lignin content in Col-0 leaves. In contrast, the lignin content in *sapx* did not show any cold-related signatures (Fig. 6). Besides, lignin content measurements from *sapx* resulted in high variation and did not provide a decent basis for further interpretations. Taken together, we concluded that cold exposure but not *B. cinerea* infection promoted plant lignification. Pathogen-triggered lignification was reported before, but mainly after inoculation with bacterial pathogens or in the response of resistant plants to otherwise necrotrophic fungi (Menden et al., 2007; Eynck et al., 2012; Lee et al., 2019; Kim et al., 2020; Jeon et al., 2023). We assume that the absence of *B. cinerea*-triggered lignification is not caused by fungal lignin degradation as it was reported earlier that *B. cinerea* is most likely not capable of degrading lignin (Hörmann et al., 2013).

**Figure 6.**
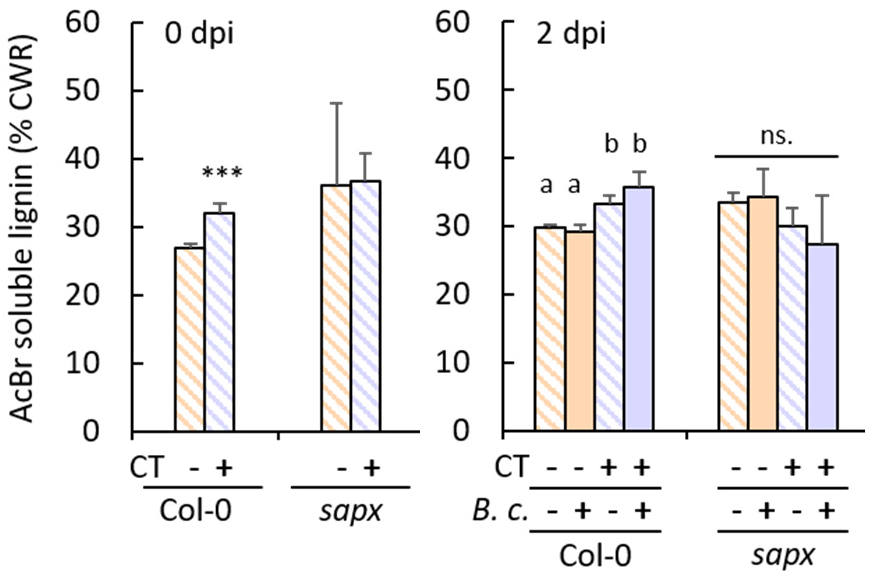
Lignin content in cell walls of leaves of Col-0 and *stromal ascorbate peroxidase* knockout (*sapx*) after prior cold pretreatment (CT +; 4 °C, 24 h) and *Botrytis cinerea* (*B*.*c*) spray infection (2 x 10^5^ spores ml^-1^ (+), Vogel buffer (-)). Bars show the mean percentage (%) of acetyl bromide-soluble lignin in the fraction of isolated cell wall residues (CWR) from three independent experiments (n = 4 – 5). Left panel: data from samples analyzed immediately after the cold exposure at the day of infection (0 dpi), two-tailed *t-test, P* ≤ 0.05 (*), 0.01 (**), 0.001 (***). Right panel: samples analyzed at 2 dpi. Different letters denote statistically significant differences within each genotype (Tukey-HSD, *P* ≤ 0.05). AcBr = acetyl bromide; CWR = cell wall residues; dpi = days post inoculation

## DISCUSSION

A single 8-24 h lasting pre-exposure to cold conditions (4°C) boosts immunity in 4-week-old Arabidopsis Col-0 against a subsequent infection with the hemibiotroph bacterial pathogen *Pst* (Wu et al., 2019; Griebel et al., 2022). Also, daily repetitive 1.5 h cold (4°C) exposures enhance plant resistance against *Pst* (Singh et al., 2014). Much less is known about the impact of cold pretreatments on plant immunity against necrotrophic pathogens. Here, we investigated the effects of preceding cold exposures on the susceptibility against the necrothrophic fungus *B. cinerea*. When *B. cinerea* (B05.10) spore inoculation was performed on cold-pretreated plants, we observed a reduction of disease lesion diameters (Fig. 1B,C). This indicates that Arabidopsis Col-0 benefits from a temporally close and timely limited cold pretreatment with an immunity boost against a broad spectrum of virulent pathogens with biotrophic and necrotrophic lifestyles. Interestingly, a milder cold exposure to only 12°C but for a longer time period of 3 days results in enhanced susceptibility of 2-week-old Arabidopsis seedlings against *B. cinerea* and *Pst* (Garcia-Molina and Pastor, 2024).

Short but more severe cold exposures and other abiotic stress treatments can be memorized in Arabidopsis for up to 5-7 days (Ding et al., 2012; Singh et al., 2014; van Buer et al., 2019). If such a memory of an experienced stress exposure results in an improved response during a subsequent stress exposure, this is defined as priming (Hilker et al., 2016). However, when cold exposure and pathogen inoculation are separated by 5 days at normal growth conditions, only plant resistance against the hemibiotroph *Pst* is potentiated (Griebel et al., 2022; Schütte et al., 2024) but not plant defence against the necrotrophic *B. cinerea* (Fig. 1). Hence, the cold priming memory is beneficial for defence against virulent hemibiotrophic *Pst* but not supportive for the defence against the necrotrophic fungus *B cinerea*. Host resistance mechanisms vary according to the lifestyle of the pathogen (Liao et al., 2022). We assume that defence responses enhanced by a cold priming memory rather include signal pathways beneficial for immunity against virulent biotrophs.

Recently, we reported that the effects of cold priming on plant resistance against bacterial pathogens relies on the availability of plastid APX (Griebel et al., 2022; Schütte et al., 2024). Here, we showed that the transient cold-enhanced resistance against *B. cinerea* immediately upon cold exposure was also stronger when plastid APX were present (Fig. 2). How do plastid APX contribute to plant pathogen defence subsequent to an abiotic stress exposure? APX scavenge ROS by using ascorbate as an electron donor (Asada, 1999). At the end of a 10-24 h lasting cold exposure (4°C), 4-week-old Arabidopsis have transiently enhanced ROS amounts (van Buer et al., 2016; Wu et al., 2019). When we measured ROS 2 h after the cold at the time point of inoculation (Fig. 4A, 0 dpi), we could not anymore observe the cold-triggered ROS burst. This indicates a fast reacclimation to a precold ROS homeostasis at normal growth conditions. Transient ROS waves are often observed in plant stress signalling (Lamb and Dixon, 1997; Apel and Hirt, 2004; Miller et al., 2009; Peláez-Vico et al., 2024). In response to a post-cold *B. cinerea* inoculation, cellular and apoplastic ROS generation was stronger in cold-pretreated than in control plants at 1 dpi. (Fig 4). In our study, higher *B. cinerea*-triggered ROS levels in cold-pretreated Col-0 plants correlated with higher resistance (Fig. 1,4). However, the contribution of plant ROS in defence against necrotrophic pathogens is complex (Torres et al., 2006). For instance, Pogány et al. (2009) suggested that ROS generated by plasma-membrane located NADPH oxidase RBOHD support plant resistance during the early phase of defence against necrotrophic fungi but enhance susceptibility at later stages. Also, transgenic *Nicotiana tabacuum* lines with less chloroplastic ROS triggered by *B. cinerea* at later stages of infections (3 dpi) showed enhanced resistance against this necrotrophic fungus (Rossi et al., 2017). We suggest that the enhanced ROS levels observed already at 1 dpi in our study of cold-pretreated Arabidopsis are connected with smaller fungal lesions and a strengthened plant resistance. Interestingly, sAPX-but not tAPX,-deficient lines responded with stronger *B. cinerea*-triggered ROS generation even without a cold-pretreatment (Fig. 4). This indicates that sAPX scavenges a significant proportion of *B. cinerea*-triggered ROS in the wildtype, but also in *tapx*, and that *tAPX* contributes less than *sAPX* to the amount of fungal-triggered plant ROS amounts. In addition, the enhanced ROS levels in *sapx* lines matched with a slightly, but not significantly, improved resistance of control *sapx* against *B. cinerea* (Fig. 3). In contrast, the weakened fungal-triggered ROS response of cold-treated *sapx* plants compared to cold-treated Col-0 or control sAPX was connected with weaker cold-enhanced resistance (Fig. 2,4). The measured ROS levels in *tapx* did not differ from Col-0. We suggest, that the stromal ROS protection with contribution from sAPX but without tAPX is sufficient to result in a wildtype-like plastid ROS homeostasis. In contrast, tAPX, but not sAPX, was identified as a specific mediator of cold priming-dependent altered activation of stress-related transcripts during a second cold exposure and affects cold priming-dependent regulation of chloroplast NADPH dehydrogenase activity (van Buer et al., 2016; van Buer et al., 2019; Seiml-Buchinger et al., 2022). Our ROS measurements (Fig.4) support the concept of distinct functions for sAPX and tAPX on plastid ROS control and redox signalling in response to cold pretreatments.

Among other functions, the interplay of apoplastic ROS and peroxidases plays pivotal roles in modifying and remodelling plant cell walls (Kärkönen and Kuchitsu, 2015). Cell wall modifications also occur in responses to abiotic stress exposures, such as cold (Le Gall et al., 2015). In biotic stress interactions, plant cell walls are physical barriers and provide protection against the invasion of pathogens (Underwood, 2012). Necrotrophic fungi secrete a large repertoire of cell wall degrading enzymes to facilitate their successful infection. To counteract this, plants sense pathogens by monitoring cell wall integrity and activate defence pathways including remodelling of cell walls (Bellincampi et al., 2014; Lee et al., 2019; Pontiggia et al., 2020; Wolf, 2022; Kim et al., 2023). Precursors of lignin derive from the phenylalanine ammonia-lyase (PAL) pathway and Arabidopsis *PAL1* is induced during cold exposure (Rohde et al., 2004; Olsen et al., 2008; van Buer et al., 2016; Griebel et al., 2022). We observed an enhanced lignification in Arabidopsis Col-0 after the 24 h lasting cold exposure (Fig. 6), which might hamper fungal penetration and support plant resistance. By contrast, degradation of lignin or additional lignification was not detectable at 2 dpi with *B. cinerea* (Fig. 6). It was already proposed by Hörmann et al. (2013) that *B. cinerea* is not able to degrade lignin. Avirulent PTI/ETI-triggering *Pst* strains and to a lower extent the virulent *Pst* strain, promote plant lignification as part of induced plant immune responses (Lee et al., 2019). Lignin enhances disease resistance against the hemibiotrophic *Pst* (Lee et al., 2019) and might also contribute to resistance against the necrotrophic fungus *B. cinerea* (Fig. 1, 6).

The most prominent plant cell wall modification in response to pathogen detection is the deposition of callose (Nishimura et al., 2003; Ellinger and Voigt, 2014). Our findings (Fig 5) corroborate previous observations (Wu et al., 2019) that cold exposure triggers callose formation in Arabidopsis leaf tissue (in the absence of pathogens). Cold-triggered callose formation was absent in *sapx*, but pathogen-triggered callose deposition did not depend on sAPX (Fig. 5). This indicates an additional and novel contribution for plastid sAPX in cold-triggered callose formation and distinguishes cold and pathogen-responsive pathways for callose deposition.

In summary, we conclude that also resistance to necrotrophic pathogens benefits from a short preceding cold treatment with increased resistance that correlates with stronger ROS formation, cold stress-induced lignification and callose deposition.

## MATERIAL AND METHODS

### Plant material and cultivation

Experiments were carried out with *Arabidopsis thaliana* accession Columbia-0 (Col-0) and described knockout lines *sapx* and *tapx* (Kangasjarvi et al., 2008). All lines are in Col-0 background. Plants were cultivated in round pots (Ø 6 cm) on a substrate composed of Topferde (Einheitserde, Germany), Pikiererde (Einheitserde, Germany), Perligran Classic (Knauf, Germany) in a 14:14:5 ratio supplemented with 0.5 g liter^-1^ dolomite lime (Deutsche Raiffeisen-Warenzentrale, Germany). After sowing, seeds were stratified at 4°C for 2 days and seedlings were pricked out approximately 8 days after stratification. Plants were grown in a controlled environmental chamber at constant humidity (60 ± 5 %) with 10 h of light (100 – 120 μmol photons m^-2^ s^-1^; Luminol Cool White fluorescence stripes, Osram, Germany) and a temperature of 20°C ± 2°C during the day and 18°C ± 2°C during the night (14 h).

### Cold stress treatments

Cold treatments were performed as previously described (van Buer et al., 2016; van Buer et al., 2019; Griebel et al., 2022). Briefly, four-week-old plants were exposed to cold 2.5 h after onset of light by transferring them to a growth chamber with a constant temperature of 4 ± 2°C but otherwise identical aeration, illumination, and air humidity as in the 20°C chamber. After a continuous cold exposure for 24 h (comprising a full day and night phase), the plants were placed back to the 20°C chamber, labelled, and randomized with the non-cold-treated control plants. After 2 h (CT, cold treatment set-up) or 5 days (CP, cold priming set-up) at normal growth conditions (20°C), plants were used for pathogen infection assays

### *Botrytis cinerea* infection

Inoculations with the *Botrytis cinerea* strain B05.10 were performed 4.5 h after onset of light. For drop inoculation assays, spore suspensions were adjusted to 5 x 10^4^ spores ml^-1^ potato dextrose broth (PDB, 6 g l^-1^). Spray-inoculation was carried out with spore suspensions adjusted to 2 x 10^5^ spores ml^-1^ in Vogel buffer (sucrose 15 g l^-1^, tri-sodium citrate 2.5 g l^-1^, K_2_HPO_4_ 5 g l^-1^, MgSO_4_ x 7 H_2_O 0.2 g l^-1^, CaCl_2_ x 2 H_2_O 0.1 g l^-1^, NH_4_NO_3_ 2 g l^-1^, pH 6). The adjusted spore suspension was incubated under gently agitation at room temperature for 4 h to allow germination. Drop inoculation was conducted by pipetting a 6 μl droplet next to the midrib of a fully expanded leaf. The mean of four drop-inoculated leaves of one plant represents one biological replicate in the lesion diameter experiments. Spray inoculations were carried out by evenly spraying the leaf surface with the spore suspension. Spore-free Vogel buffer was used for mock spray treatments as control. Infected plants were kept as described above except for an additional high humidity environment created by watering and light permeable coverage of the pots.

### Quantitative real-time PCR analysis

Transcript analyses were performed using quantitative real-time PCR (qRT-PCR). One sample (= one biological replicate) consisted of 4 pooled plant rosettes. Plant samples were harvested at the indicated time points. RNA extractions, cDNA syntheses and qRT-PCR assays from ground plant material were performed as previously described (van Buer et al., 2016; Griebel et al., 2022). A mix of oligo(dT)16 primers and random primers was used for cDNA syntheses. The qTower^3^ G instrument (Analytik Jena, Germany) was used for qRT-PCR assays. Primers for genes of interest and reference genes are listed in Table 1. Values of cycle thresholds were determined using qPCRsoft (Analytik Jena, Germany) and relative expression of transcripts of interest were calculated using the ΔCT-method against the geometric mean of two reference genes *Yellow Leaf Specific Protein 8* (*YLS8*) and *RGS1-HXK1 Interacting Protein 1* (*RHIP1*).

**Table 1.**
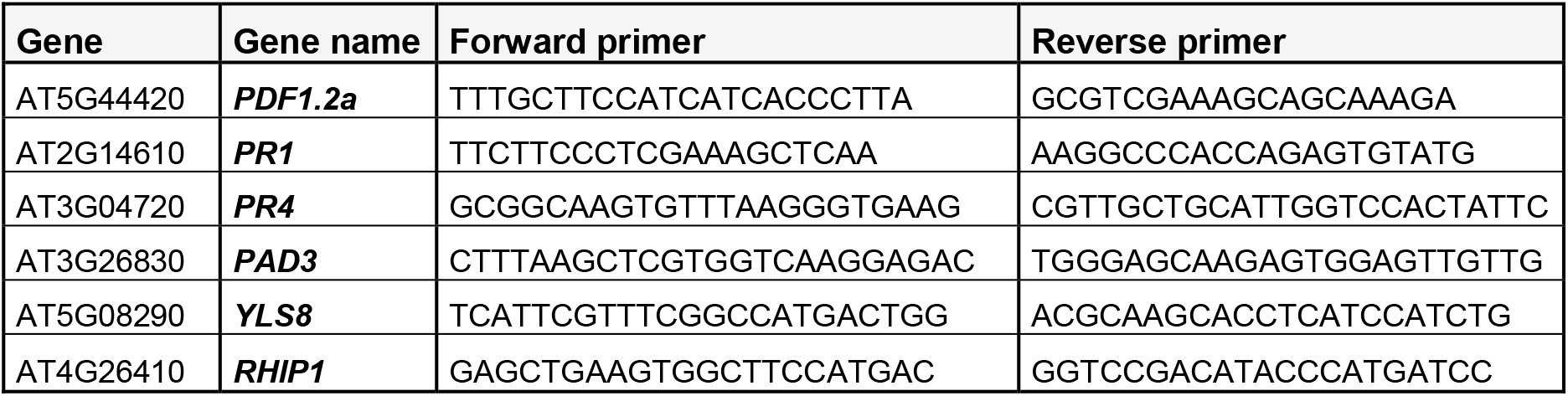
Primers used in this study.

### DAB-staining

To analyse hydrogen peroxide abundance after cold exposure and *B. cinerea* infection, 3,3′-diaminobenzidine (DAB) was used as previously described (Bittner et al., 2020). 1 mg ml^-1^ DAB was dissolved in phosphate-buffered saline (PBS; 73 mM NaCl, 10 mM, Na_2_HPO_4_, 3 mM KCl, 2 mM KH_2_PO_4,_ pH 7.2). A minimum of 4 leaves from 3 plants per treatment was harvested and collected in a tube with DAB-staining solution. The collected leaves were infiltrated with the staining solution by applying a gentle vacuum for 5 min in a desiccator. Afterward, samples were incubated overnight in the dark. The background of stained leaves was removed in a 1:1:3 mixture of acetic acid, glycerol, and ethanol before analysis and image acquisition. The intensity of DAB staining per leaf was calculated using ImageJ (Schneider et al., 2012; Bittner et al., 2020).

### Luminol-based ROS assay

ROS was measured by horseradish peroxidase (HRP)-catalyzed luminol oxidation in the presence of H_2_O_2_. 21 h after *B. cinerea* spray-inoculation, leaf discs (Ø 4 mm) from at least 3 plants per treatment were collected in _dd_H_2_0-filled petri dishes and incubated for 3 h under normal growth conditions to minimize wounding response during the measurement. Thereupon, leaf discs were transferred to 96-well plates. 150 μl reaction solution (200 μM luminol, 0.04 μg HRP ml^-1^ _dd_H_2_0) was added to each well. Chemiluminescence was measured with a CLARIOstar^PLUS^ plate reader (BMG Labtech, Germany) and is shown as the sum of all counted relative light units (RLU) within 1 h of measurement. RLU of inoculated samples were normalized to the corresponding mean of the non-infected mock treatments (RLU_treatment_ – RLU_mock treatment_ = normalized RLU) within each independent experiment.

### Callose quantification

Callose was stained with aniline blue (Carl Roth GmbH, Germany) according to Schenk and Schikora (2015) with the alteration of an extended aniline blue incubation overnight. One biological replicate of the callose quantification was defined as the mean of five randomly imaged field of views (7 mm^2^) from one leaf. Leaves were taken from three different plants and the experiments were repeated three times independently. Monochrome pictures were taken using a AxioImager Z2 microscope (Zeiss, Germany) equipped with a Zeiss filter 49 (G

365/ FT 395 / BP 445/ 50) and an Axiocam 712mono (Zeiss, Germany) camera. Images were acquired with the software ZEN blue (Zeiss, version 3.3) and numbers of callose depositions were manually counted.

### Lignin quantification

Quantifications of lignin were performed according to Moreira-Vilar et al.(2014) with modification from Chezem et al. (2017). 14 leaf discs (Ø 8 mm) from at least 3 plants were pooled to one sample and defined as one biological replicate. Samples were frozen in liquid nitrogen and ground in 2 ml tubes, each containing 2 x 4 mm glass beads, using a swing mill (Retsch, Germany) at 30 Hz for 2 min. Ground samples were vacuum-dried using a Concentrator plus (Thermofisher, Germany) and the dried leaves were ground again as described above. Next, cell wall residues (CWR) were twice extracted in an ultrasonic bath with each of the following solvents for 15 min: 1 ml methanol, 1 ml PBS with 0.1 % (v/v) Tween 20, 1 ml ethanol, 1 ml chloroform/methanol (1:1 ratio) and at least 1 ml acetone. After each extraction step, samples were centrifuged at 16.000 g for 10 min and the supernatant was discarded. CWR were dried at 45°C using a Concentrator plus and ground with a swing mill (2 ml tubes, each containing 2 x 4 mm glass beads) at 30 Hz for 10 min. Approximately 3 mg of CWR from each sample was used for the spectrometric analysis. For this purpose, 500 μl of 25 % (v/v) acetylbromid in glacial acetic acid was added to each sample and incubated at 50°C with rotation (800 rpm) for 2 h. After dissolving the lignin, samples were cooled on ice and centrifuged at 20.000 g for 15 min. 125 μl of the lignin-containing supernatant was mixed with 500 μl glacial acetic acid and 250 μl of 5 M hydroxylamine HCl / 2 M NaOH (1:9 ratio). Photometric measurements were done in a single glass cuvette (10 mm) using an Ultrospec 2100 pro at 280 nm and a sample without CWR as blank. Acetylbromid soluble lignin content was calculated with the extinction coefficient ε of 23.35 mg cm^-1^ l^-1^ (Chang et al., 2008).

### Statistical analyses and boxplot design

The statistical analysis was conducted using Excel for Student’s t-tests, basic R environment for ANOVA and the follow-up Tukey-HSD test, the R agricolae package for LSD-Fisher test after prior ANOVA. Box plots of the summarized data were generated using the R package ggplot2 and show the median, the distance between the upper quartile (qn = 0.75) and lower quartiles (qn = 0.25), and the values of each data point as dots. Data points form independent experiments are shown in different colour.

## ACKNOWLEDGEMENTS

We would like to thank Marcel Wiermer and Philipp Rohmann (FU Berlin) for providing the *Botrytis cinerea* strain and methodical expertise on culturing the strain. We also thank Mitja Remus-Emsermann and his group (FU Berlin) for sharing access to their microscope and plate reader facilities and for providing technical support.

